# Circulation of *Klebsiella pneumoniae* strains between the termites, chimpanzees and humans

**DOI:** 10.1101/2024.10.14.618212

**Authors:** Cheikh Tidiane Houmenou, Habibou Sarr, Seydina M. Diene, Idir Kacel, Cheikh Sokhna, Florence Fenollar, Oleg Mediannikov

**Affiliations:** Aix-Marseille Univ, AP-HM, MEPHI, Marseille, France; IHU-Méditerranée Infection, Marseille, France; Aix-Marseille Univ, SSA, AP-HM, RITMES, Marseille, France; EMR MIMES, Campus International IRD-UCAD Hann, Dakar, Senegal; IRD, Marseille, France

**Author notes:** **Correspondence** : Oleg MEDIANNIKOV, **Address**: MEPHI, IHU-Méditerranée Infection; 19-21 Boulevard Jean Moulin 13385 Marseille cedex 5 France. **Institutional Review Board Statement:** The study was conducted in accordance with the Declaration of Helsinki and approved by the National Health Research Ethics Committee of the Ministry of Health and Social Action of Senegal (No.: 0000045/MSAR/CNERS/SP).

**Keywords:** Antibiotic resistance genes, *Klebsiella pneumoniae*, carbapenemases

## Abstract

The emergence and multiplication of multi-resistant microbes is becoming a major concern in medicine and scientific research, requiring an integrated, global approach to prevent their spread and develop new therapeutic solutions*. Klebsiella pneumoniae* can cause a variety of infections. It has several distinctive features, such as extended-spectrum beta-lactamase (ESBL) production and frequency in hospital-acquired infections, which set it apart from other bacteria as *Pseudomonas aeruginosa* or staphylococci and make it of particular concern in the clinical setting. In this study, we worked on four strains of multidrug-resistant *K. pneumoniae* isolated from different patients at Fann Hospital in Dakar, Senegal. Whole genomes were compared them with those of nine *K. pneumoniae* isolates from chimpanzees (CHZ) and six from termites (T_NSC) from our previous study. The comparison revealed two major genetic groups: three human strains were virtually identical to those found in three chimpanzees and four genomes extracted from termites regularly consumed by chimpanzees were identical to five *K. pneumoniae* strains from chimpanzees. In silico, we identified the resistance genes and their corresponding families from each isolate, then genotyped them before comparing them with other isolates stored in the Institute Pasteur database. These strains have never been previously identified in Senegal.

It has been shown here that multiresistant *K. pneumoniae* strains circulate between chimpanzees and termites and between chimpanzees and humans. Furthermore, these strains, often associated with nosocomial infections, have been identified and genotyped outside Senegal. Although similar strains have been already identified in tropical countries and in different intensive care units throughout the world, we hypothesize that they may originate from termite-chimpanzee ecosystem.

## INTRODUCTION

Antibiotics and antibiotics resistance are both natural occurences and have been around since the dawn of time (D’Costa *et al*. 2011). The presence of naturally multi-resistant bacteria has been demonstrated in natural reservoirs dating back to over 30,000 years ago (D’Costa *et al*. 2011), probably due to horizontal transfers or genetic mutations (SNPs, insertions or deletions) or even inherited (Olaitan, Morand, et Rolain 2014). It is now widely accepted that the origin of antibiotic resistance is the environmental microbiota (Martínez 2012). Numerous cases of antibiotic resistance genes have been discovered in environments that seemingly lack antibiotics. Several explanations can account for this phenomenon (Davies et Davies 2010). Resistance genes could be part of a pre-existing natural gene pool, such as *erm* and *van* genes in soil microbiota, or *qnr* genes in aquatic microbiota (Aminov et Mackie 2007). Another explanation involves the strong selective pressure from past antibiotic use, leading to the integration of resistance genes into bacterial genomes, accompanied by compensatory mutations that mitigate the fitness cost. Furthermore, recent studies have shown that in certain tropical African countries, or in regions remote from human activity where antibiotic pressure is very rare or even virtually non-existent, such as in the natural reserves of protected forests in Senegal or Kenya, naturally multi-resistant bacteria are found in wild animals such as chimpanzees (Kempf *et al*. 2012; Baron *et al*. 2021).

However, the effect of industrial antibiotics seems to influence the natural selection of resistant bacteria (Galán *et al*. 2013). With the advent of industrial antibiotics, some bacteria seem to have acquired one or more antibiotic resistances under drug pressure (Olaitan, Morand, et Rolain 2014). It has often been stated that the obvious evolutionary factor contributing to the dissemination of antibiotic resistance genes is the ever-increasing production and consumption of antibiotics for various purposes, from treatment to the questionable practice of feeding them to food-production animals at subtherapeutic levels for growth promotion (Aminov et Mackie 2007; Van Boeckel *et al*. 2019).

Indeed, the massive use of antibiotics seems to accelerate the selection and emergence of new resistant bacteria due to the strong pressure from antibiotic molecules that bacteria with resistant or multi-resistant genes are subjected to. This phenomenon was observed for instance in hospitals within the intensive care unit, in countries with high population density and high consumption of antibiotics such as India and Bangladesh. This is also observed in countries such as China, Indonesia or South Africa where there is an overuse of antibiotics in domestic or edible animals which are exposed to repeated small doses of antibiotics (Van *et al*. 2020).

Overall, intrinsic mechanisms of the selection, circulation and acquisition of antibiotic resistance under natural conditions are poorly studied. Currently in modern medicine, beta-lactam antibiotics, which are broad-spectrum antibiotics such as carbapenems, are used in the treatment of many serious infectious diseases. However, recent studies have shown that beta-lactam resistance is not directly caused by the excessive use of manufactured beta-lactams (Collignon *et al*. 2018; Olaitan, Morand, et Rolain 2014; Baron *et al*. 2021). Carbapenems can destroy the structure of the cell wall of Gram-stain negative bacteria and are efficient against many pathogenic bacteria (Nicolau 2008). However, there are bacterial strains resistant to carbapenems including some *Klebsiella pneumoniae* strains that produce carbapenemases which inhibit the effect of carbapenems. As a result, the spread of carbapenemase-producing strains of *K. pneumoniae* has become a serious problem and the treatment of infections caused by these pathogens is a major challenge for clinicians. Moreover, several studies have shown that *K. pneumoniae* can be an opportunistic pathogen responsible for a significant proportion of nosocomial and community-acquired infections. Indeed, *K. pneumoniae* is an important cause of pneumonia and lung abscesses as well as central nervous system infections such as meningitis and community-acquired brain abscesses, urinary tract infections and other diseases (Paczosa et Mecsas 2016). In addition to being pathogenic, this genotypic variation of *K. pneumoniae* is a major obstacle to controlling the public health risk associated with this pathogen.

A previous study (Baron *et al*. 2021) in west Africa in a protected natural reserve (Dindefello Natural Reserve) remote from major industrial human activity (factories, hospitals, human waste, big animal farms), revealed the presence of several potentially pathogenic multidrug-resistant strains of *K. pneumoniae* in the stools of chimpanzees and in the digestive tract of termites. It has been hypothesized that multidrug-resistant strains of *K. pneumoniae* that colonized chimpanzee’s intestines may originate from eaten insects. Genomic sequences of *K. pneumoniae* isolates, sometimes known to be both multidrug-resistant in humans and pathogenic in chimpanzees, have been deposited on the European Bioinformatics Institute (EBI) and made accessible to all.

In this article, we performed comparative analyses of clinical strains of *K. pneumoniae* isolated from four patients at the Fann Hospital, Dakar, Senegal, with strains previously isolated from chimpanzees and termites living in the Kedougou forest in Senegal as well as other with other human and animal strains of *K. pneumoniae* obtained from the database of the Institut Pasteur, Paris, France. For that purpose, we assessed the clusterization of these strains and searched for all the resistance genes in our clinical isolates that could justify the phenotypic aspect of their resistance to antibiotics (notably carbapenems and NDMs). Finally, these strains were genotyped in silico to identify them and compare them to other strains from Senegal, Africa and the rest of the world, which could not necessarily be sequenced entirely (amplification of specific segments of DNA that contain genetic markers specific to a single species or the use of the MLST method which is a technique that focuses on a set of housekeeping genes conserved among bacteria, generally 5 to 7 genes). These comparative analyses made it possible to detect groups of isolates that could imply a possible circulation of K. pneumoniae strains between humans, chimpanzees and termites.

## MATERIALS AND METHODS

### Source of data

Clinical data were obtained by taking samples from several patients, at different levels of intervention at the FANN hospital in Dakar, explaining the origin of the biological data (Table 1). However, the human clinical data, the isolates selection, bacteriological and molecular analyses of *K. pneumoniae* strains have been described previously (Sarr *et al*. 2023). Among the strains of enterobacteria collected from patients in Dakar, we randomly selected four clinical strains of *K. pneumoniae*. All strains were isolated from hospitalized patients in the neurology, neurosurgery, infectiology and otolaryngology departments but the selected four strains were isolated from the latter in a diagnostic laboratory as a part of bacteriological analyzes of routine (Sarr *et al*. 2023). Isolates were identified using matrix-assisted laser desorption ionization time-of-flight (MALDI-TOF) mass spectrometry (Bruker Daltonik, Bremen, Germany) (Seng *et al*. 2009). In addition, genomic data on *K. pneumoniae* from termites and chimpanzees were retrieved from the NCBI database, deposited by Baron *et al*. (Baron *et al*. 2021). For further analyses, we selected 10 *K. pneumoniae* strains isolated from chimpanzees and six strains isolated from termites described in the manuscript by Baron *et al*.(Baron et al. 2021)

**Table 1.** Origin of clinical data, extraction and collection of biological samples from Fann Hospital in Dakar, Senegal.

| Date | New Code | Hospital | Samples | Age / Sex | Ward | MALDI-TOF MS |
| --- | --- | --- | --- | --- | --- | --- |
| 07/19/2022 | MAR157 | Dakar (Fann Hospital) | Pus | 37 years old / F | Cardiovascular thoracic surgery | *K. pneumoniae* |
| 07/13/2022 | MAR163 | Dakar (Fann Hospital) | Pleural fluid | 61 years old / M | Pneumology | *K. pneumoniae* |
| 07/12/2022 | MAR168 | Dakar (Fann Hospital) | Blood culture | 15 years old / M | Cardiovascular thoracic surgery | *K. pneumoniae* |
| 07/07/2022 | MAR172 | Dakar (Fann Hospital) | Blood culture | Unknown / M | Resuscitation | *K. pneumoniae* |

### Antibiotic resistance of strains

All clinical isolates from the Fann hospital in Dakar were analyzed at the IHU-Méditerranée Infection research laboratory in Marseille, France. Antibiotic sensitivity tests were carried out using the disk diffusion method as indicated in the recommendations of EUCAST 2022 (Sarr *et al*. 2023) . Antibiograms for other strains of *K. pneumoniae* from chimpanzees and termites were extracted from the bacteriology analysis by Baron *et al*. Subsequently, we produced a table (**Table 2**) summarizing the antibiogram targeting only the antibiotics ertapenem, cefotaxim and ciprofloxacin on the *K. pneumoniae* from chimpanzees, humans and termites to compare the sensitivity state (the bacterial vulnerability) of all strains.

**Table 2.** Tests the sensitivity/resistance of *K. pneumoniae* strains described here and in the manuscript of Baron *et al*. to three antibiotics (ertapenem, cefotaxim and ciprofloxacin)

| Isolat strains of *Klebsiella pneumoniae* |  |  |  |  |  |  |  |  |  |  |  |  |  |  |  |  |  |  |  |  |
| --- | --- | --- | --- | --- | --- | --- | --- | --- | --- | --- | --- | --- | --- | --- | --- | --- | --- | --- | --- | --- |
| Hosts | Chimpanzees (CHZ) |  |  |  |  |  |  |  |  |  | Termites (NCS) |  |  |  |  |  | Human (MAR) |  |  |  |
| Strains |  |  |  |  |  |  |  |  |  |  | NCS | NCS | NCS | NCS | NCS | NCS | MAR | MAR | MAR | MAR |
| Antibiotics | CHZ1 | CHZ2 | CHZ3 | CHZ5 | CHZ6 | CHZ7 | CHZ16 | CHZ17 | CHZ30 | CHZ35 | 35A | 43A | 43B | 45A | 45B | 50D | 157 | 163 | 168 | 172 |
| ETP | - | - | - | + | + | - | + | + | - | + | - | - | - | - | - | - | + | - | + | + |
| CTX | + | + | + | + | + | + | + |  | + |  | + | + | + | + | + | - | - | - | - | - |
| CIP | + | + | + | + | + | - | + | - | - | - | - | - | - | - | - | - | + | + | + | + |

Legend: abbreviation and coding of antibiotics:
+ Resistance (by red color) CHZ = from chimpanzees MAR = from human - Sensitive (by green color) NCS = from termites
Table representing the sensitivity/resistance profile of isolated *K. pneumoniae* strains regarding three antibiotics: Ertapene (ETP, Beta lactam family), Cefotaxim (CTX, Beta-lactam family) and Ciprofloxacin (CIP, Fluoroquinolone family)

### Extraction and sequencing of bacterial isolates

#### DNA extraction

Genomic DNA (gDNA) of bacteria was extracted as follows: After resuspending strains in 160 µl of G2 Buffer (Digestion Buffer from Qiagen-EZ1 DNA Tissue Kit), a mechanical lysis with the FastPrep-24™ 5G Grinder instrument (mpBio) at maximum speed (6.5 m/sec) for 90 seconds was performed followed by an enzymatic lysis using lysozyme (40µl per sample Sigma) incubation at 37°C for 30 minutes.

DNA was extracted on the EZ1 advanced XL biorobot (Qiagen) with EZ1 DNA tissue kit. The elution volume was 50 µL. gDNA was quantified by a Qubit assay with the dsDNA High Sensitivity Assay kit (Thermofisher Scientific).

#### Sequencing

In this article, we used the MiSeq sequencer of *K. pneumoniae*. Genomic DNA was sequenced on the MiSeq technology instrument (Illumina Inc, San Diego, CA, USA) with the paired end strategy and was barcoded in order to be mixed with others genomic projects prepared with the “Nextera XT Library Prep Kit” (Illumina). To prepare the paired end library, dilution was performed to obtain 1 ng of each gDNA as input (0.2 ng/µl in 5 µl). The tagmentation step fragmented and tagged the DNA. Then limited cycle PCR amplification (12 cycles) completed the tag adapters and introduced dual-index barcodes.

After purification with AMPure XP beads (Beckman Coulter Inc, Fullerton, CA, USA), the libraries were then normalized on specific beads according to the Nextera XT protocol (Illumina). Normalized libraries were pooled for sequencing on the MiSeq. Automated cluster generation and paired end sequencing with dual index reads were performed in a single 39-hours run in 2x250-bp with MiSeq reagent Kit (V2-500 cycles) (Illumina). The paired reads were filtered according to the read qualities. The raw data were configured in fastq files for R1 and R2 reads. The number of reads generated for each sample with their corresponding species is summarized in **Table 3**.

**Table 3.** The number of reads and genes generated by each sample after sequencing and assembly.

| Name samples on Miseq | Number of reads | Name MO query | Assembly tool | Number of contigs | Genome size (bp) | % GC | Genes number | CDS number |
| --- | --- | --- | --- | --- | --- | --- | --- | --- |
| MAR157 | 2485982 | *K. pneumoniae* | Spades | 95 | 5,851,204 | 56.6 | 5544 | 5457 |
| MAR163 | 944146 | *K. pneumoniae* | Spades | 149 | 5,576,559 | 57.1 | 5275 | 5189 |
| MAR168 | 1335426 | *K. pneumoniae* | Spades | 134 | 5,846,238 | 56.6 | 5549 | 5462 |
| MAR172 | 1031122 | *K. pneumoniae* | Spades | 137 | 5,846,839 | 56.5 | 5534 | 5447 |

### Genome assembly

After sequencing we used the SPAdes script to assemble the reads of each sequenced strain. SPAdes (St. Petersburg genome assembler) Genome Assembler is an open-source tool for de novo sequencing. It is an assembly toolkit containing various assembly pipelines. The current version of SPAdes works with Illumina or IonTorrent reads and is capable of providing hybrid assemblies using PacBio, Oxford Nanopore and Sanger reads. You can also provide additional contigs that will be used as long reads (Prjibelski *et al*. 2020). To see more details on the installation and the mode of use, I invite you to go to the following site https://github.com/ablab/spades. SPAdes require a 64-bit Linux system or Mac OS and Python (supported versions are Python 2.7, and Python3: 3.2 and higher) to be pre-installed. After sequecing the reads output are obtained is in FastQ format. To launch the assembly of a klebsiella genome we used basic options of SPAdes on the Linux terminal command line, we started by calling the python script of the Spades tool (spades.py: the main executable script) on a terminal, then fill in the two fastq files (read 1 and read 2: left or right paired reads) resulting from the sequencing on the same command line. Also, enter the --only-assembler (runs assembly module only) option to do simple assembly and finally give the name of the output file (is the name given to the assembly results folder) containing the fasta sequence. The output folder will contain a fasta file (scaffold.fasta) consisting of multiple contigs and/or scaffolds. After assembly we will move on to cleaning scaffolds since some of them have a small size (less than 800bp) and others have a very low coverage rate (Calculate the average coverage of all scaffolds then remove anything that is less than 25% of the average coverage rate). Then we renamed each genome file by encoding by combining each Klebsiella pneumoniae strain with the host organism and their antibiotic resistance phenotype (**Supplementary Table 1**).

### Whole genome comparison

OrthoANI (Orthologous Average Nucleotide Identity) shows the similarity values between two genome sequences. This is an improved version of ANI (Average Nucleotide Identity) that complies with OGRI rules. (Organism-specific Genomic Islands) (Lee *et al*. 2016). It can be used for classification and identification of bacteria with a suggested proposed cutoff for species delineation of 95∼96%.

Later, a faster version was developed, named OrthoANIu, using usearch program instead of BLAST. Although ANI is widely used to classify and identify bacteria, OrthoANI was developed to overcome the large differences in reciprocal ANI values associated with the ANI algorithm (Yoon *et al*. 2017). Furthermore, OrthoANIu tool employs USEARCH over BLAST for its OrthoANI calculations which increases the number of comparative studies and substantially decrease computational time. The software tools are available as web-service and standalone program. The OrthoANI tool can be used to compare two genomic sequences and define a percentage of identity between the two sequences.

However, in this project, we used OrthoANIu to compare several strains of *Klebsiella* genomes, namely our four clinical strains of *K. pneumoniae* (MAR157, MAR163, MAR168 and MAR172); two clinical isolates (*K. pneumoniae* strain_Kp3412_LBHALD and *K. pneumoniae* strain_Kp3739_LBHALD) from another project by Kempf et al. (Kempf et al. 2012); six *K. pneumoniae* isolates from termites; nine *K. pneumoniae* isolates from chimpanzees from the study by Baron et al. and three *Klebsiella africana* isolates retrieved from the NCBI database (with the following accession numbers: CP084874, CP059391 and JAQCON01000001) to form the outgroup of the phylogenetic tree.

To achieve this comparison, the tool compares the sequence strains two by two and gives the percentage of identity each time, thus generating a matrix of genomic identity percentages. Using the orthoANI command line tool first requires installing Java Runtime Environment Version 8 (Java Download) and the USEARCH (<u>usearch11.0.667_i86linux32.gz</u>, It is compatible with 32-bit).

Although 64-bit version will reduce overall process time, we concluded that it is sufficient to use 32-bit version for a small number of genomes. On the command line, you will need to fill in the folder containing all the fasta files of the genomes to be compared, fill in the output of the comparison result in a matrix format with the -fmt matrix option, then give the name of the output file containing the matrix of two-by-two comparison of genomes. To do this, usearch executes the orthoANI software in a Java environment to generate the comparison result in the form of a matrix.

We developed a Rstudio script (compatible with Rtudio version 3.5) to generate a genome strain comparison heatmap. The program receives a well-determined data matrix and returns a heatmap containing the more or less identical strains according to the percentage of identity detected by the OrthoANI tool.

### Generation of phylogenetic trees and clusters

We used scapper (https://github.com/tseemann/scapper) to align the genomes of *K. pneumoniae* isolated from chimpanzees (CHZ), termites and humans (Dakar patients as well as including the genomes described by the study by Kempf *et al).* The genomes of three isolates of *K. africana* were used as an outgroup for the tree (Figure 2). The distance between isolates was calculated using the iqtree 2 tool (Minh *et al*. 2020) with boostrap, which is an efficient and versatile phylogenomic software using the maximum likelihood model (http://www.iqtree.org) then visualized by itol (https://itol.embl.de/) on the online website interactively (Letunic and Bork 2024).

**Figure 1:**
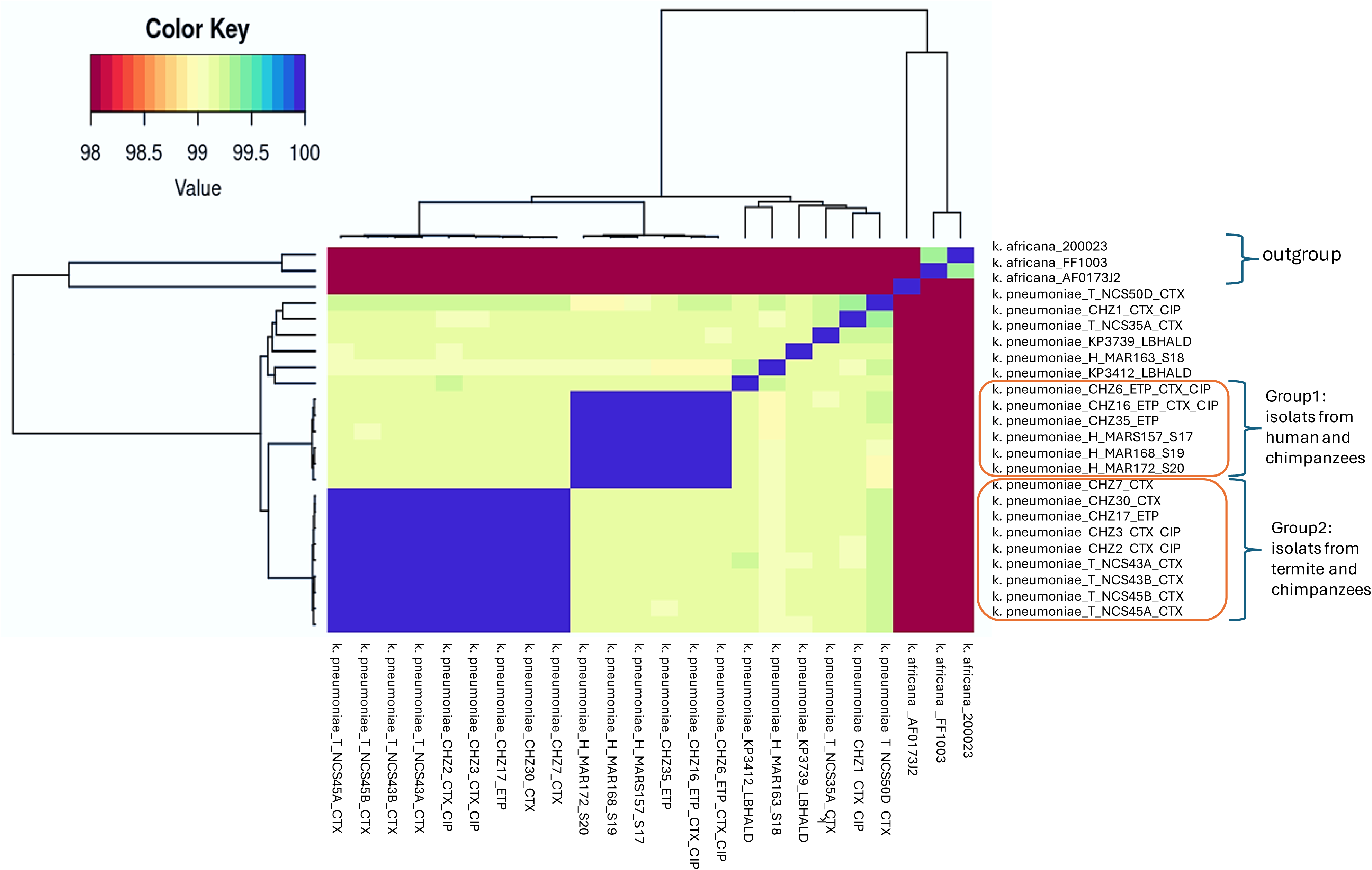
Genomic comparison of different strains of *K. pneumoniae* via the orthoAni. **T** : Termite ; **CHZ** : Chimpanzee ; **H** : Human; **ETP** : Ertapenem resistant ; **CTX** : Cefotaxim resistant; **CIP** : Ciprofloxacin resistant; **ETP**_CIP : resistant to Ertapenem and Ciprofloxacin; **ETP_CTX_CIP** : resistant to three antibiotics

**Figure 2.**
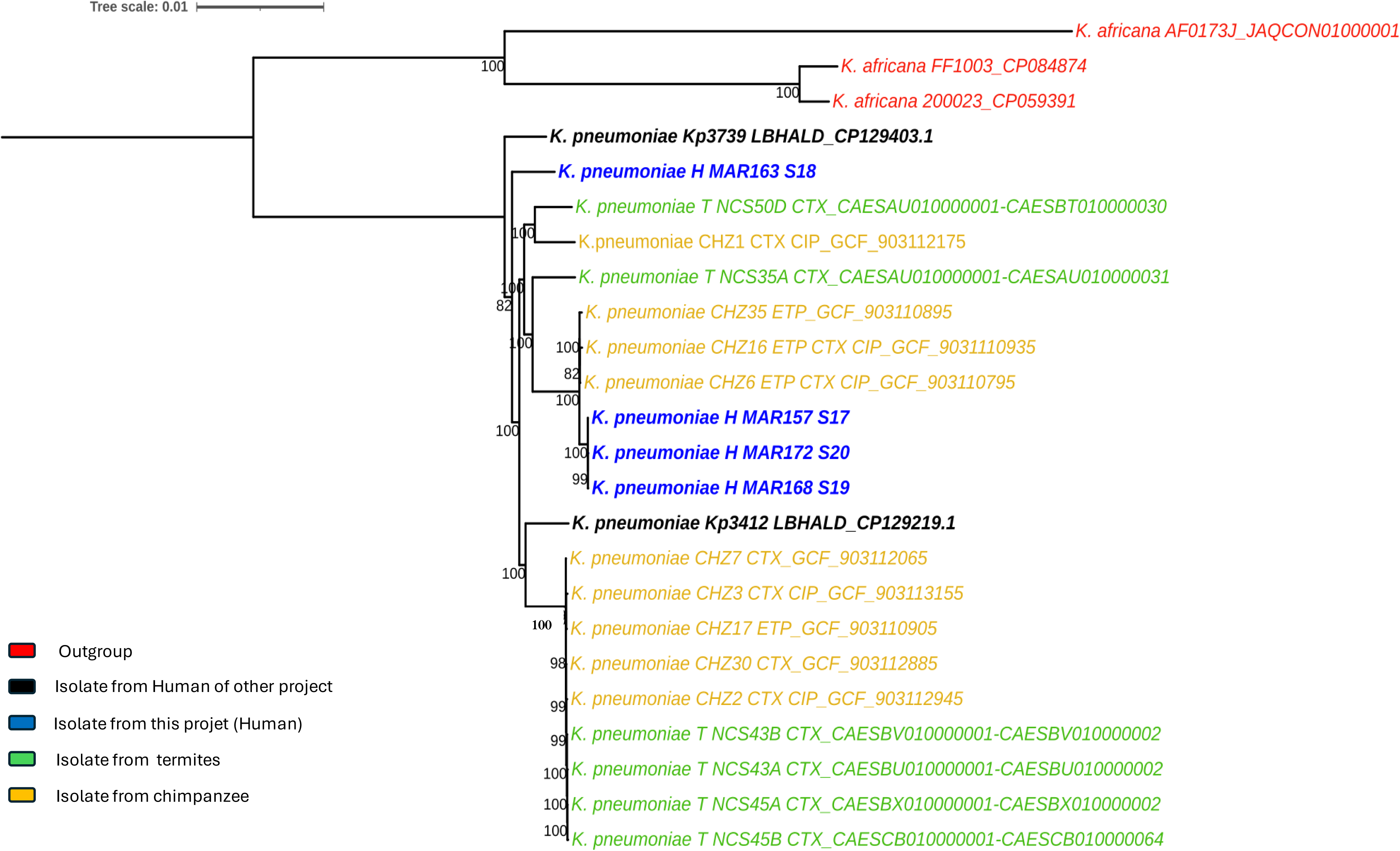
Phylogenetic tree of different *K. pneumoniae* genome isolates from chimpanzee, termite and human organisms in comparison with *K. africana.* Tree interfered using the maximumliklihood model, with a bootstrap value of 1000.

### Genotypic definitions of *K. pneumoniae* isolates based on MLSTs

The BIGSdb-Pasteur web platform is an integrated resource for genomics-based strain taxonomy, offering reference nomenclature databases, genomic libraries, and comparative genomics tools. The platform is powered by the Bacterial Isolate Genome Sequence database software (University of Oxford). BIGSdb-Pasteur hosts a collection of multilocus typings (MLST), full genome typings (core genome cgMLST) and additional patterns (antimicrobial resistance, virulence, …) for bacterial pathogens of public health importance, including *K. pneumoniae* and *Listeria monocytogenes*.

Contextual, genomic and typing data for thousands of bacterial isolates are also accessible and searchable via the website (https://bigsdb.pasteur.fr/). First, we identified the typical pattern of each of our four strains isolated from patients in the database (BIGSdb-Pasteur) which is based on the search for seven typical determining genes (*gapA*, *infB*, *mdh*, *pgi*, *phoE*, *rpoB* and *tonB*) of *K. pneumoniae*. After obtaining the sequence types (ST), we then compared them with the *K. pneumoniae* ST’s from Senegal, Africa and finally with those from the rest of the world. We used the MLST tool (Seemann T, mlst Github https://github.com/tseemann/mlst) to confirm our *in silico* genotyping results, by analyzing the contigs of each genome against the traditional PubMLST typing schemes (https://pubmlst.org/) which is a collection of organized open-access databases that integrates population sequence data with provenance and phenotypic information for over 130 different microbial species and genera (Jolley, Bray, et Maiden 2018) (**Table 5**).

### Identification of antibiotic resistance genes

ABRicate (Seemann, T. Abricate. GitHub Repository. https://github.com/tseemann/abricate) is a mass contig screening tool for antimicrobial resistance or virulence genes. It comes with several databases including NCBI, CARD, ARG-ANNOT, Resfinder, MEGARES, EcoOH, PlasmidFinder, Ecoli_VF and VFDB. In our case, we only used one database ARG-ANNOT (Gupta *et al*. 2014) because it is more compatible with our analysis by providing supplementary necessary information regarding new existing and putative antibiotic resistance genes, resistance gene family membership, broad beta-lactamase spectrum genes. In this project, we used ABRIcate to identify all the resistance genes contained in the different strains of *K. pneumoniae* in this study and to categorize the pathogenicity of these strains. Moreover, ABRicate enables us to find out which resistance genes these strains of *K. pneumoniae* had in common.

## RESULTS

### Origin of clinical data

For further analyses, we have selected four antibiotic-resistant strains from previously analyzed collections of enterobacteria strains from two Senegalese hospitals (Sarr *et al*.). All four strains were isolated from clinical samples of patients hospitalized in the cardiovascular thoracic surgery, cardiovascular surgery, pneumology and intensive care departments of the Fann Hospital in Dakar, Senegal (**Table 1**).

One strain (MAR157) was obtained from the pus of a 37-year-old female patient whereas the second (MAR163) was isolated from the pleural fluid of a 61-year-old man. Strain MAR168 was isolated from the blood of a 15-year-old patient, hospitalized in the cardiovascular department. Finally, the MAR172 strain was obtained by blood culture from a man hospitalized in the intensive care unit . The four strains, MAR157, MAR163, MAR168 and MAR172, isolated from these patients were identified using MALDI-TOF MS as *K. pneumoniae* with high identity scores of 2.40, 2.03, 2.32 and 2.11, respectively.

### Antibiotics resistance of clinical strains

Using several antibiogram tests, the strains were evaluated for antibiotic resistance and the results showed different profiles depending on the type of resistance (ertapenem, cefotaxime and ciprofloxacin). All strains were found to be sensitive to cefotaxim (CTX) but resistant to ciprofloxacin (CIP). Notably, only the MAR163 isolate is sensitive to ertapenem (ETP). The clinical data results were cross-referenced with those from antibiotic testing work by S. Baron *et al*. to compare the susceptibility of *K. pneumoniae* from chimpanzees, termites and humans (**Table 2**).

When compared with *K. pneumoniae* strains CHZ5, CHZ6 and CHZ16 isolated from wild chimpanzee’s stools and termite strains from southern Senegal (**Table 2**), we found evident similarities. Chimpanzee’s *K. pneumoniae* were resistant to the three antibiotics while the NCS50D strain from *Macrotermes* termites (common protein source consumed by chimpanzees) was the only sensitive to all the tested antibiotics.

### Genome sequencing and assembly

After sequencing of four clinical strains, different numbers of reads were obtained for each clinical isolate. We noticed that we have slightly fewer reads with the MAR163 isolate compared to others (**Table 3**). However, once assembled using the SPades assembler tool, genomes had approximately the same sizes, around 5.7 Mbp.

Additionally, 95 contigs were obtained with the isolate MAR157 while the rest of the strains resulted in contigs numbers ranging from 134 to 149. Moreover, there were fewer genes in strain MAR163 (5275 genes) compared to the others. In fact, the number of genes for the other strains comprised between 5534 and 5560 genes. Despite these differences, the blast result of four strains clearly identified them as *K. pneumoniae*.

### Genome comparison and phylogenetic analysis

The raw result of the orthoANIu calculation is presented in the form of a matrix, which can be transformed into a comparative heatmap. The heatmap comparison shows two significant clusters: a first cluster of isolates bringing together *K. pneumoniae* from chimpanzees and termites and a second cluster consisting of chimpanzee and human strains. Interestingly, two human clinical isolates in Senegal from a recent study (*K. pneumoniae*) were not included within the two clusters detected. The phylogenetic tree also highlighted two clusters of strains similarly as the orthoANI results. The two clinical strains in the study by Kempf *et al*. were not found in either cluster, and their phylogenetic relationships were more distant from those of both clusters. There results confirm those obtained using orthoANI.

### Resistome analysis of investigated *K. pneumoniae* isolates

The use of the ABRicate tool allowed the detection of several resistance genes targeting a large panel of antibiotics (**Table 4**).

**Table 4.**
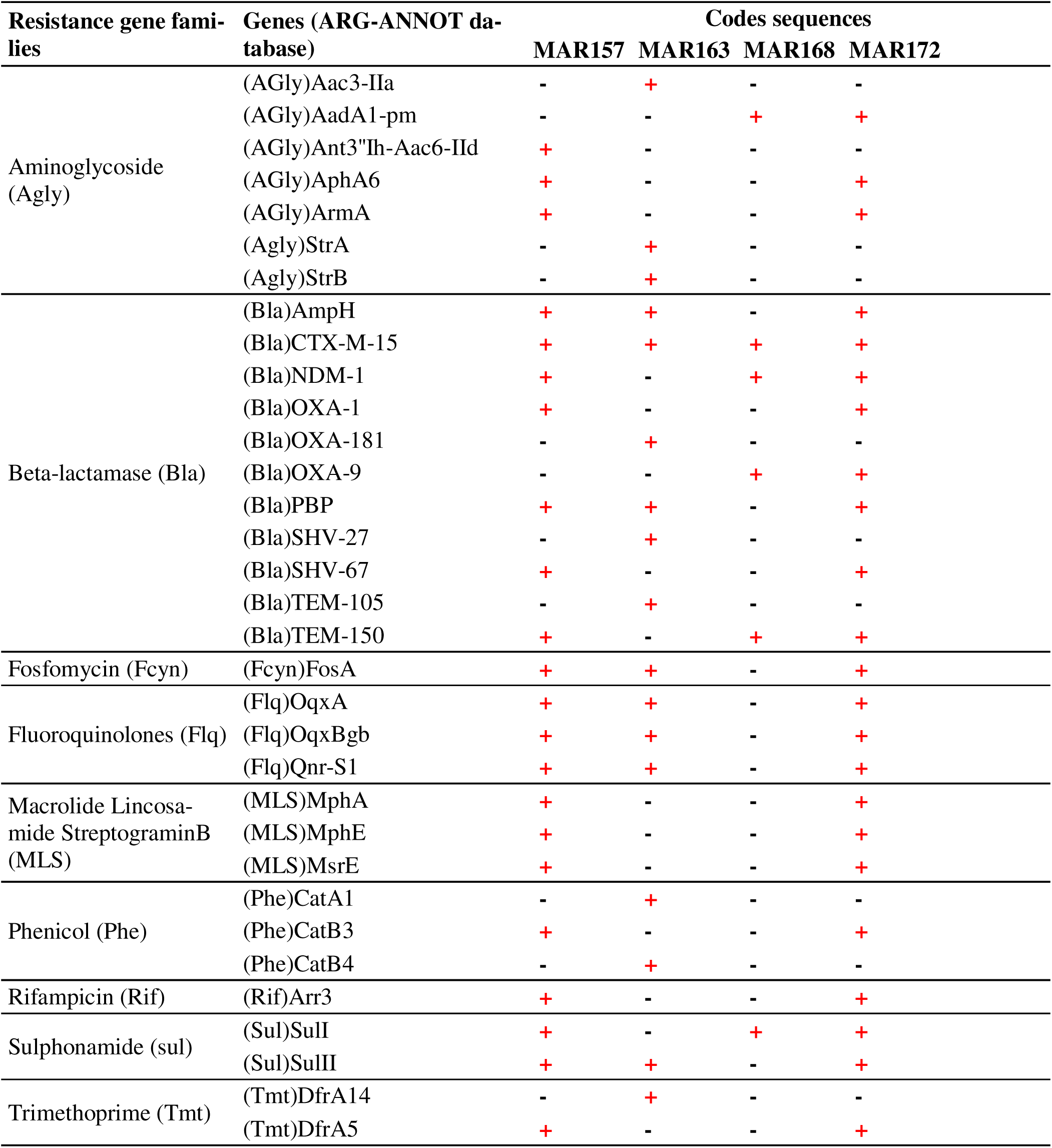
Detection of resistance genes in clinical strains.

The investigation of antibiotic resistance genes (ARG) in the four sequenced *K. pneumoniae* isolates using the ABRicate pipeline revealed the presence of genes conferring resistance to several classes of antibiotics including aminoglycosides (n = 7), β-lactams (n = 11), fosfomycin (n = 1), fluoroquinolones (n = 3), macrolide-lincosamide-streptogramin B (n = 3), phenicols (n = 3), rifampicin (n = 1), sulphonamide (n = 2), and trimethoprime (n = 2) (**Table 4**). In detail, apart from the strain MAR168 which exhibits only six ARGs including three β-lactam resistance genes (*bla*_CTX-M-15_, *bla*_NDM-1_, *bla*_OXA-9_, and *bla*_TEM-150_), the three other isolates (i.e. MAR157, MAR163, and MAR172 isolates) harbored 22, 17, and 23 ARGs respectively, which confers resistance to all tested antibiotic classes (**Table 4**). Indeed, aminoglycoside resistance genes such as *aphA*6, *armA*, or *aadA*1 were identified in these isolates; and genes of interest agaisnt β-lactams including the pandemic metallo-β-lactamase *bla*_NDM-1_ and Extended-spectrum-β-lactamase (ESBL) *bla*_CTX-M-15_ genes were detected in these three isolates (**Table 4).** Other genes including fosfomycin resistance gene fosA, quinolone resistance gene qnrS1, sulphonamide resistance gene sul2, fluoroquinolone resistance gene Flq, sulfonamide resistance gene sulI and beta-lactamase resistance gene (Bla)-PBP were detected in three out of four isolates namely MAR157, MAR163 and MAR172. Interestingly, the ESBL *bla*_CTX-M-15_ gene was detected in all four *K. pneumoniae* isolates investigated in this study suggesting a dissemination of this gene within the hospital.

### Genotypic definitions of *K. pneumoniae* isolates based on MLSTs

The *in silico* genotyping of our four isolated strains identified two distinct sequence types (STs), 147 (ST147) and 234 (ST234) (**Table 5**). Three (MAR157, MAR168 and MAR172) of the four human strains were identified as ST147 and only one strain MAR163 had a typical sequence identified as ST234. When compared with other termite and chimpanzee strains, we found that they were quite different from the others except three chimpanzee strains (CHZ35, CHZ6 and CHZ16). Indeed, these three chimpanzee strains have the same ST147 sequence type as those found in three patients of the Fann hospital in Dakar. In addition, by comparing them with the two other *K. pneumoniae* genomes (K_pneumoniae_Kp3412_LBHALD and K_pneumoniae_Kp3739_LBHALD) recently sequenced in Senegal from the study by Kempf et al. (Kempf et al. 2012) we note that our clinical strains have STs (ST147, ST234) different from these two strains (ST39 and ST133, Table 5). Furthermore, our strains were compared with 179 *K. pneumoniae* strains isolated in Senegal that are genotyped and deposited in the Pasteur site’s bigsdb database, none of them had the same ST (ST147 and ST234) as our patients’ four strains (Supplementary Table 2). Comparison with termite and chimpanzee strains (Table 5), showed a very different ST’s, except for the 3 chimpanzee strains (CHZ35, CHZ6 and CHZ16).

**Table 5.**
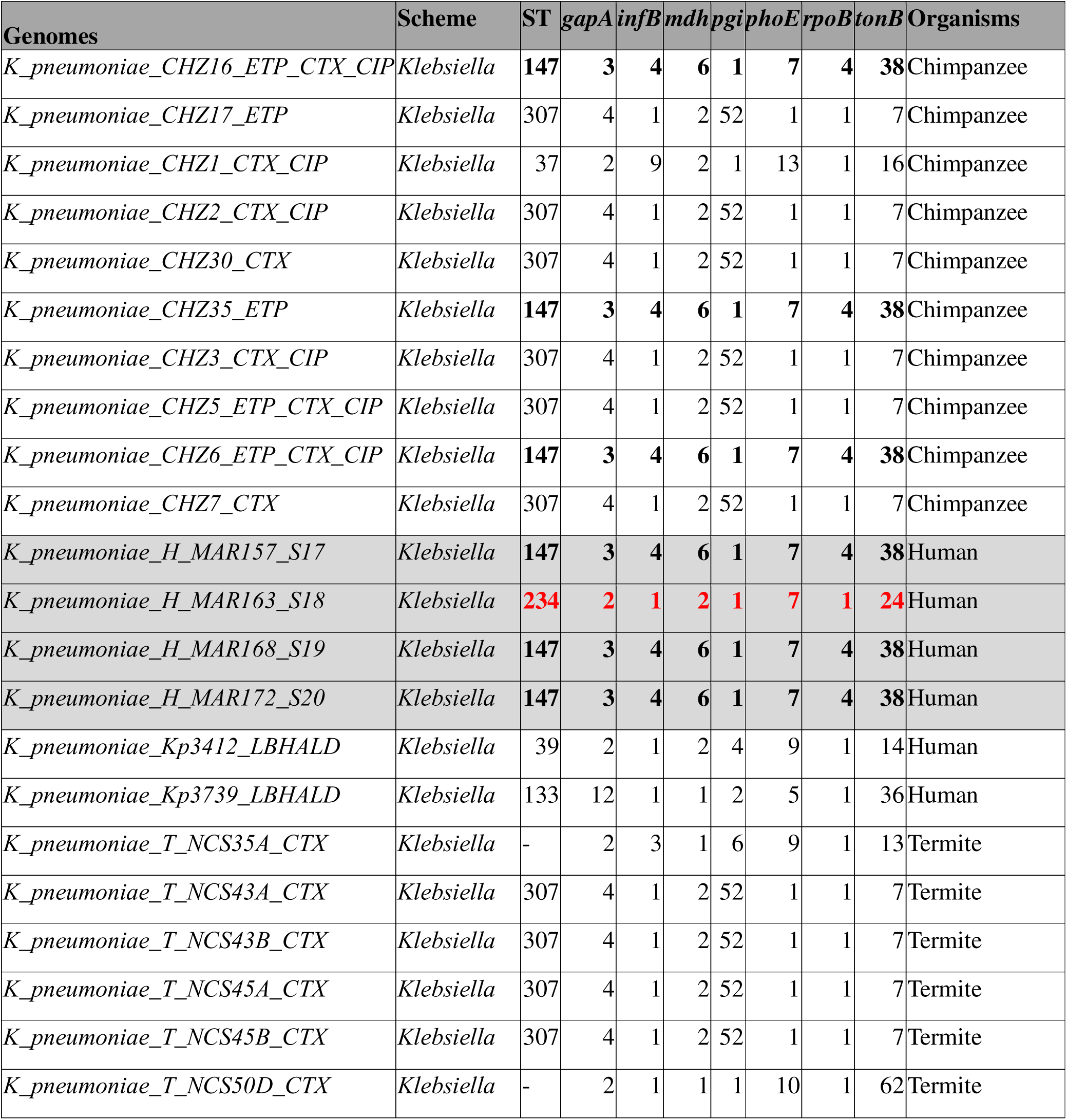
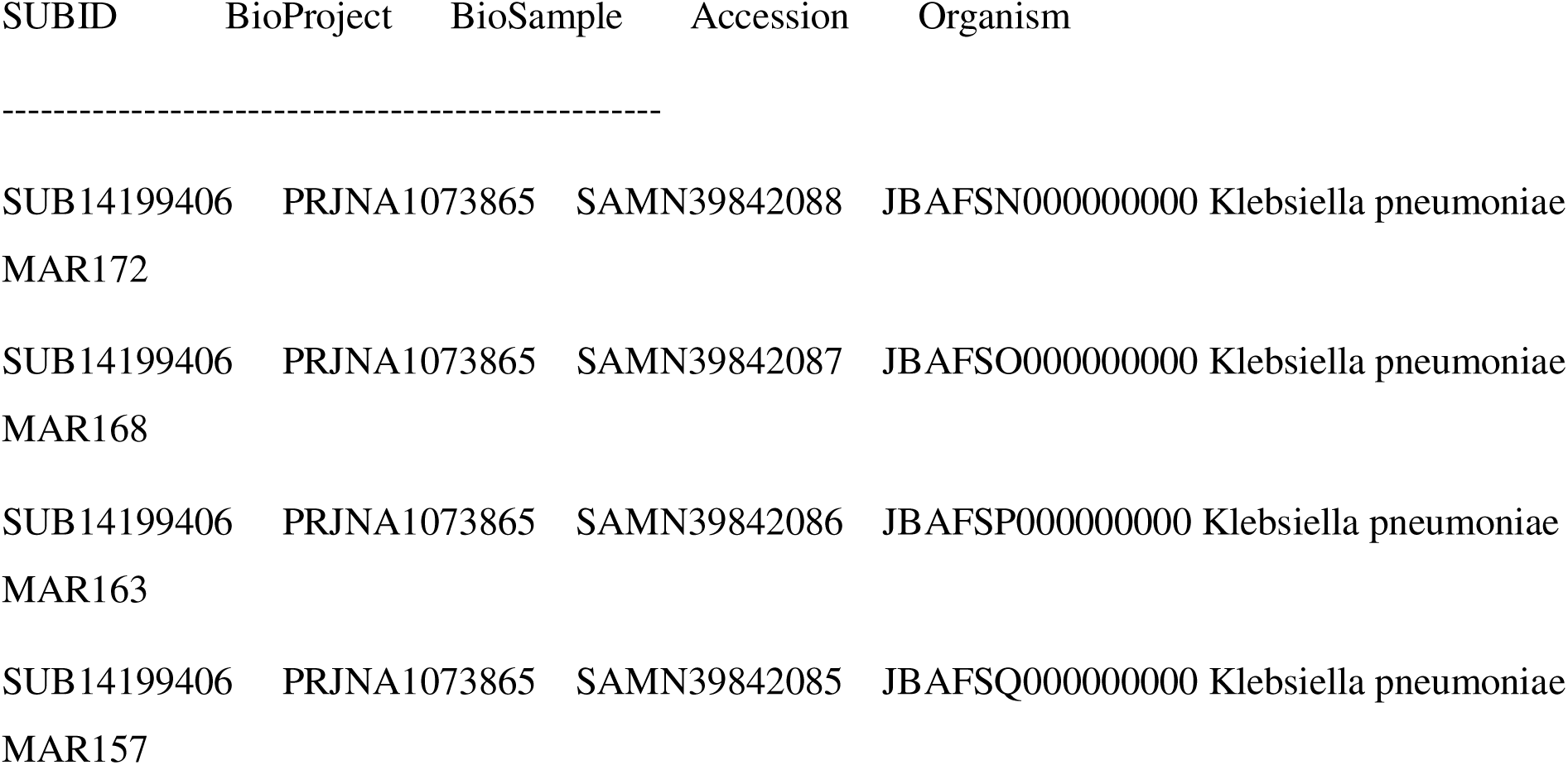
*In silico* genotyping: Analyzing contig files against traditional PubMLST typing schemes.

Indeed, these three *K. pneumoniae* from chimpanzee showed the same typical patterns and sequences as those found in patients from Dakar. We then compared the genotyping results with those of two other *K. pneumoniae* genomes (strain Kp3412_LBHALD and strain Kp3739_LBHALD) from humans of another sequencing project, to see if our strains have a similar profile to those found in our Dakar patients. Interestingly, our four strains were identified as genotypes present in countries other than Senegal (Figure 3: a and b).

**Figure 3.**
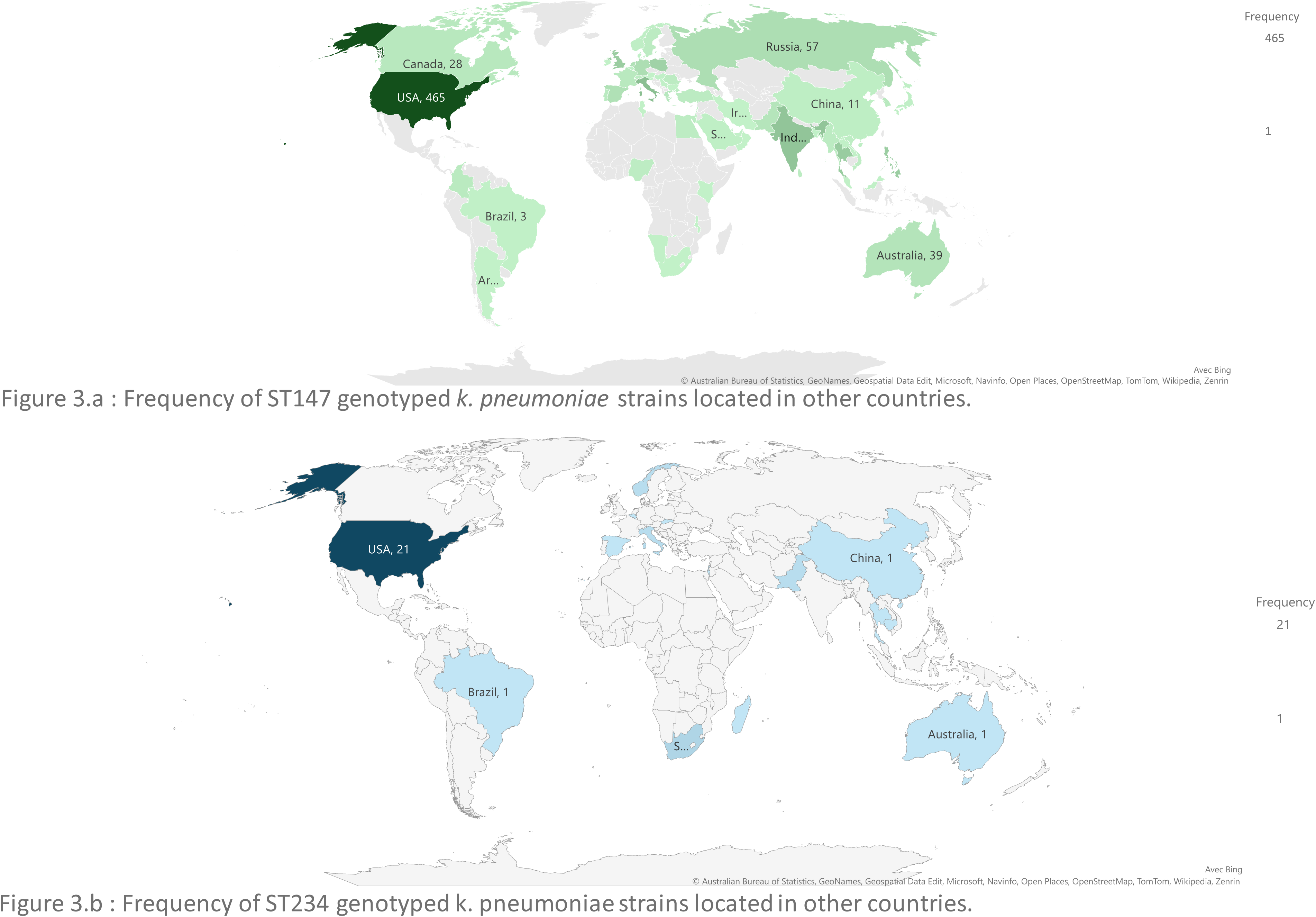
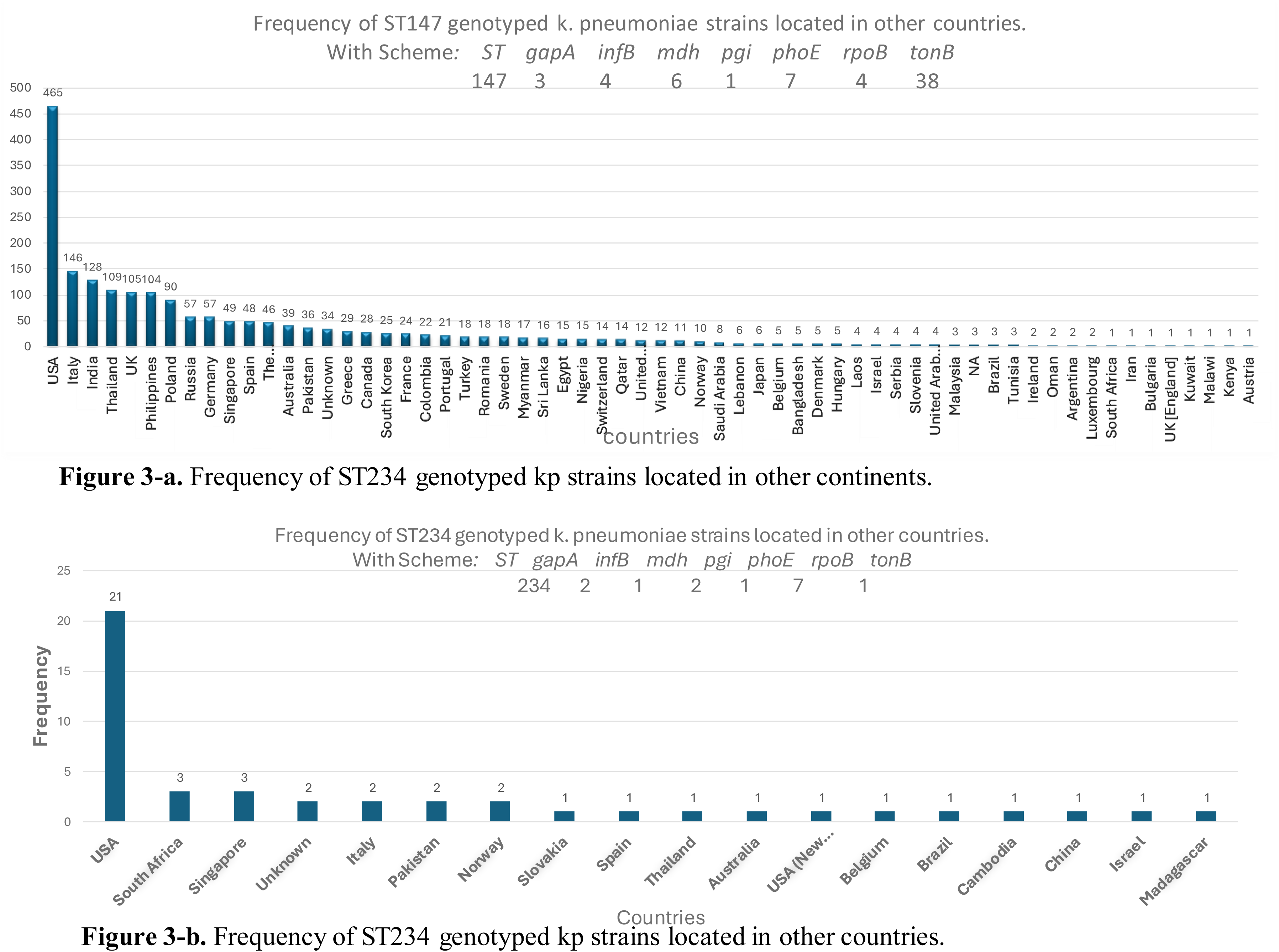
**a.** Frequency of ST147 genotyped k. pneumoniae strains located in other countries. **b.** Frequency of ST234 genotyped k. pneumoniae strains located in other countries.

## DISCUSSION

Worldwide, the epidemiology of antibiotic resistance genes is becoming a serious threat to public health (Nakacubo Gitta, Wabwire-Mangen, et Tshimanga 2011). In low- and middle-income countries, such as in sub-Saharan Africa and west Africa more specifically, well-equipped laboratories are rare, and the means used to fight the growth of antimicrobial resistance are very weak. Therefore, we are witnessing the spread of antimicrobials and their residues into the environment, contributing to the selection of resistant bacterial strains, including human and animal pathogens (Kagambèga *et al*. 2024). In Senegal, a study on the epidemiology of resistance genes in enterobacteria revealed several isolates of multi-resistant bacteria from patients hospitalized at the Fann hospital in Dakar (Sarr *et al*. 2023). Among them, four were identified as *K. pneumoniae*. Another study carried out in Kedougou in a protected forest in the south of Senegal, far from any habitable zone and any human activity, revealed the existence of multi-antibiotic-resistant *K. pneumoniae* in chimpanzees stool and in the digestive tract of termites (Baron *et al*. 2021). Interestingly, our study revealed that these isolates (four clinical strains, 10 isolates of *K. pneumoniae* from chimpanzees and six isolates of *K. pneumoniae* from termites) resulted into two well-defined cluster groups according to the orthoANI analysis and confirmed by phylogenetic tree and *in silico* genotyping The first cluster consisting of isolates from chimpanzees and termites shows almost perfect identity between them and proves the hypothesis which was introduced by the study by Baron *et al*. (Baron *et al*. 2021), the contamination of chimpanzees through the consumption of termites. Additionally, the second cluster, composed of chimpanzee isolates and clinical strains, shows an almost identical resemblance between the last two, except for the MAR163 isolate which is excluded from the cluster, and not belonging to any group. This is the first proof of circulation between its three hosts.

Genome analysis showed that three of the four *K. pneumoniae* strains **(Figure 1)** isolated from humans (nosocomial infections at Dakar hospital) were identical to previously isolated chimpanzee strains. This is the second evidence of the circulation of *K. pneumoniae* ESBL strains in the termite/chimpanzee/human ecological niche. The results obtained on the phylogenetic tree (Figure 2) reconfirmed the two clusters already detected in the heatmap from the orthANIu results (Figure 1).

There could be several explanations for this possible circulation of *K. pneumoniae*. Such as, the common consumption of termites (infected with bacteria in their digestive tract) by chimpanzees in forests, the exceptional use of chimpanzees as a meal for humans, knowing that these species are highly protected in Kedougou by the forest rangers. However, the most plausible hypothesis is the use of water from a natural source (lake, pond, river, etc.) shared by humans and chimpanzees in villages, or even the contamination of shared food from tree fruits or leaves. A study carried out in sub-Saharan Africa, in Burkina Faso more precisely in the surroundings of the city of Ouagadougou, showed that 36% of the bacteria collected (out of 264 samples collected) were multi-resistant K. pneumoniae and producers of beta-lactamase (ESBL-Kp). Its ESBL-Kp were prevalent in stormwater runoff and treated and untreated wastewater.(Kagambega *et al*. 2024). All of these avenues remain to be explored. In contrast, the two K. pneumoniae clinical isolates Kp3412_LBHALD and Kp3739_LBHALD from Kempf et al. were isolated and did not group with our Dakar clinical strains, although they were collected at the Bacteriology laboratory of CHU Aristide Le Dantec in Dakar, Senegal.

Furthermore, mining of the BIGSdb-Pasteur database showed more than 46,173 Klebsiella isolates recorded worldwide including 6,222 K. pneumoniae genotyped with sequence types identified and classified based on Multilocus Sequence Typing (MLST). This database allowed us to study in depth and in a more advanced manner the distribution and circulation of strains by comparing on the one hand our isolated strains with each other and on the other hand by comparing them with isolates from Senegal, from Africa and the rest of the world by providing a global picture of the genotypic status of K. pneumoniae.Indeed, several species of bacteria have been isolated in Senegal and in other African countries including *K. pneumoniae* (Shinga Wembulua *et al*. 2021) but have never been the subject of genomic extracts and sequencing for more in-depth analyses. This can become a limitation to perform genomic comparisons of strains or to use new techniques for describing new species based on bioinformatic analysis recourses such as orthoANI, DDH *in silico* or even phylogenetic tree reconstruction. Genomic comparison results from genotyping our clinical isolates with strains from termites, chimpanzees and clinical data from Kempf et al. show that isolates with the same sequence type cluster together (Table 4). In our case, strains MAR157, MAR168 and MAR172 from clinical data from Fann Hospital, Dakar, having the same ST147 as CHZ6, CHZ16 and CHZ35 strains from chimpanzees (Table 4), clustered together (Figure1 et Figure 2). De même les isolats issus de termites (NCS43A, NCS43B, NCS45A et NCS45B) et de chimpanzés (CHZ2, CHZ3, CHZ7, CHZ17, et CHZ30) ayant le même ST307 (Tableau 4) se clustérisent ensemble (Figure 1 et Figure 2). These results not only provide a genotypic explanation for the two clusters detected in the orthoANI (Figure 1) and phylogenetic tree (Figure 2) results, but also demonstrate that isolates from the same host type (e.g. chimpanzees) can group together in different clusters (chimpanzee-human and chimpanzee-termite) due to their ST difference (CHZ with ST147 and CHZ with ST307). Strain MAR163 does not clustardize with either our clinical strains from Fann Hospital or those collected from Le Dentec Hospital in Dakar from the work of Kempf et al. This may be explained by their genotypic difference. Isolate MAR136 has an ST234 that is different from those of our three other strains (ST147) and Kempf’s two isolates (ST133, ST39). Our explanations also show that isolate MAR163 to ST234 has a pattern composed of alleles from 7 loci (gapA, infB, mdh, pgi, phoE, rpoB and tonB) and that it has only two loci (pgi, phoE) with the same allele numbers (1 and 7 respectively) in common with our other 3 strains (MAR157, MAR168 and MAR172). This genetic difference at the other 5 loci (gapA, infB, mdh, rpoB and tonB) may be explained by mutation or insertion/selection. His new findings could be evidence of natural antibiotic resistance in bacteria (D’Costa et al. 2011), despite the presence of an anti-microbial effect. We also sought to determine the extent of circulation of ST147 and ST234 K. pneumoniae throughout Senegal and the rest of the world, and to compare them with our clinical strains from the Fann Hospital in Dakar. Of the 179 K. pneumoniae isolates genotyped in Senegal, none had a typical sequence at ST147 or ST234. This information shows just how genetically different K. pneumoniae isolates can be, but also that these strains are either new isolates in Senegal, or we don’t have enough data (bacteria, K. pneumoniae) genotyped or sequenced. On the other hand, ST147 is found in six African countries (Egypt, Nigeria, Tunisia, South Africa, Malawi and Keynan) and ST234 only in South Africa and Madagascar (Figure 3: a and b). Circulation of multidrug-resistant K. pneumoniae has been demonstrated between chimpanzees and humans, and between termites and chimpanzees, but no isolate clusters have been found between humans and termites, thus limiting the almost perfect circulation of this resistant bacterium between its three hosts (humans-chimpanzees-temites). This could be explained by the paucity of clinical data collected at Fann Hospital and the small number of termite isolates (six strains) compared with those from humans (four strains).

For instance, ST147 was detected in 62 countries, with a higher frequency of genotyping in countries such as the USA (465 ST147 isolates), Italy (146 isolates), India (128) or Thailand (109) than in other countries such as Saudi Arabia (8 isolates), the United Kingdom (5 isolates) or Kenya (1 isolate) (Figure 3.a). For ST234, only 18 countries including the USA (21 isolates), South Africa (3 isolates), Singapore (3 isolates) and Italy (2 isolates) have the ST234 genotype (Figure 3.b). This difference in the frequency of ST147 and ST234 between Africa and the rest of the world could be explained either by the low circulation of these strains in northern countries, or by the lack of biological data due to poverty factors (lack of resources, lack of laboratory facilities). This high frequency in developed countries (USA, Italy, UK) could be explained by the number of health facilities (hospitals, health posts) where surveillance systems, such as those of the European Union (EARS-Net) and the USA (CDC), actively monitor the prevalence of antibiotic resistance.

Understanding resistance genes and how they spread is crucial to slow the rise in antibiotic-resistant strains and preserve the effectiveness of treatments (CDC 2024). Carbapenems, such as ertapenem, are often considered antibiotics of last resort to treat severe multidrug-resistant (MDR) *Klebsiella pneumoniae* infections. *K. pneumoniae* can develop resistance mechanisms to carbapenems, such as the production of carbapenemases (e.g. KPC: Klebsiella pneumoniae Carbapenemase, NDM: New Delhi Metallo-β-lactamase), representing a major threat to public health. However, this bacterium can also present resistance mechanisms to fluoroquinolones, such as mutations in genes coding for topoisomerases or efflux pumps, leading to reduced sensitivity to this drug, or even produce enzymes called extended-spectrum beta-lactamases (ESBL), which are capable of degrading cephalosporins such as cefotaxime. The study of the resistance to ertapenem, ciprofloxacin and cefotaxime in *Klebsiella pneumoniae* is essential, as these antibiotics represent key classes in the treatment of bacterial infections (Pitout et Laupland 2008; Davies et Davies 2010). The 20 strains included in our study showed different resistance profiles for three antibiotics: ertapenem (ETP, belonging to the Carbapenem antibiotic family), ciprofloxacin (CIP, belonging to the Fluoroquinolone antibiotic family) and cefotaxim (CTX, belonging to the 3rd generation Cephalosporin family of antibiotics). Among isolates from chimpanzees, three (CHZ5, CHZ6 and CHZ16) out of 10 are completely resistant to the three antibiotics and out of six isolates from termites only one (NCS50D) is sensitive to all three. By comparing the antibiograms of clinical isolates against the same antibiotics, only strain MAR163 has a profile different from the others. This can be explained by the existence of resistance genes in some and their absence in others, probably due to the presence or absence of plasmids with different resistance genes in some strains. Alternatively, it may simply be a strain with a different genetic profile from the other three clinical strains(Gogoi et al. 2023; Zhu et al. 2023).

Results from resistance gene detection show that our four strains have (Bla)CTX-M-15, which are enzymes of the beta-lactamase family. It is noteworthy that CTX-M has emerged as an important cause of systemic infectious syndromes, such as community-acquired pneumonia and urinary tract infections (Pitout et Laupland 2008). As described in the results section, MAR163 has a lower gene number and a unique ST compared to the other clinical isolates. Interestingly, MAR163 was sensitive to all three antibiotics which can be explained by the absence of the resistance genes that usually detected in plasmids (Levy et Marshall 2004). Conversely, the (Bla)NDM-1 gene was detected in all three clinical strains (MAR157, MAR168 and MAR172). Studies have shown that NDM is a metallo-beta-lactamase (MBL) which uses metal ions (often zinc) to hydrolyze carbapenems and other beta-lactams. NDM has spread rapidly throughout the world, causing epidemics in many hospitals. It can be transferred via plasmids, facilitating transmission between different bacterial species. *K. pneumoniae* can become extremely resistant to antibiotics thanks to NDM, making it one of the main causes of difficult-to-treat nosocomial infections (Nordmann, Naas, et Poirel 2011; Tzouvelekis et al. 2012).

Although better equipped to manage infections, hospitals in developed countries remain vulnerable to nosocomial epidemics of resistant strains, particularly in intensive care units. This is in contrast to the situation in developing countries, where limited resources for hospital infection control, lack of access to quality healthcare and overcrowded infrastructures are thought to favor the spread of multi-resistant bacteria such as Klebsiella pneumoniae (Nordmann, Naas, et Poirel 2011; CDC 2024). According to Pasteur’s Big database, resistant strains of Klebsiella pneumoniae, particularly those producing carbapenemases such as NDM and KPC, are spreading worldwide. Although NDM was first identified in India, it has spread to hospitals all over the world, including in developed countries. Globalization and international travel play a key role in this spread.

## Conclusion

Antibiotic resistance is a serious problem that is of great concern to all public health and scientific research. This study allowed us to know the dimension of genetic variability within the same species but also allows us to better understand that the same species of bacteria (*K. pneumoniae*) can be found in several different organisms (termites, chimpanzees, humans). Except for the MAR163 isolate, the other three strains contained the NDM-1 resistance gene (metallo-β-lactamase from New Delhi) while this same gene was found in *Escherichia coli* and it is capable of breaking down several important antibiotics, including penicillin’s, cephalosporins and carbapenems (Qamar *et al*. 2023).

Furthermore, a study focusing on the notable spread of carbapenem-resistant *K. pneumoniae* (Ahmed El-Domany *et al*. 2021) revealed that the NDM-1 gene was identified among the genes found in *K. pneumoniae* isolates, knowing that, *K. pneumoniae* resistant to carbapenems constitutes a serious threat to human health. In our article, we demonstrated a possible circulation of *K. pneumoniae* among its three organism groups. The passage of this bacterium from termites to chimpanzees can be explained by the latter’s consumption of termites, but we do not yet know its mechanism of transmission from chimpanzees to humans.

