## supplementary data for "Circulation of *Klebsiella pneumoniae* strains between the termites, chimpanzees and humans"

**Supplementary Table 1**. Encoding strain names: we rename each strain of *K. pneumoniae* by giving the name of the organism of origin from which it was extracted and then linking it with the result of the antibiotic sensitivity test.

| **Number** | **Strain genome** | **Organism** |
| --- | --- | --- |
| 1 | K_pneumoniae_CHZ1_CTX_CIP.fna | Chimpanzee |
| 2 | K_pneumoniae_CHZ2_CTX_CIP.fna | Chimpanzee |
| 3 | K_pneumoniae_CHZ5_ETP_CTX_CIP.fna | Chimpanzee |
| 4 | K_pneumoniae_CHZ6_ETP_CTX_CIP.fna | Chimpanzee |
| 5 | K_pneumoniae_CHZ17_ETP.fna | Chimpanzee |
| 6 | K_pneumoniae_CHZ7_CTX.fna | Chimpanzee |
| 7 | K_pneumoniae_CHZ3_CTX_CIP.fna | Chimpanzee |
| 8 | K_pneumoniae_CHZ30_CTX.fna | Chimpanzee |
| 9 | K_pneumoniae_CHZ3_CTX_CIP.fna | Chimpanzee |
| 10 | K_pneumoniae_CHZ35_ETP.fna | Chimpanzee |
| 11 | K_pneumoniae_H_MAR157_S17.fasta | Human |
| 12 | K_pneumoniae_H_MAR163_S18.fasta | Human |
| 13 | K_pneumoniae_H_MAR168_S19.fasta | Human |
| 14 | K_pneumoniae_H_MAR172_S20.fasta | Human |
| 15 | K_pneumoniae_T_NCS43B_CTX.fna | Termite |
| 16 | K_pneumoniae_T_NCS35A_CTX.fna | Termite |
| 17 | K_pneumoniae_T_NCS45A_CTX.fna | Termite |
| 18 | K_pneumoniae_T_NCS45B_CTX.fna | Termite |
| 19 | K_pneumoniae_T_NCS50D_CTX.fna | Termite |
| 20 | K_pneumoniae_T_NCS43A_CTX.fna | Termite |

The name of each strain was renamed by encoding the name with the organism of origin and the initials of the antibiotics it resists.

**Supplementary Table 2.** Table summarizing the 179 genotyped kp strains, classified by sequence type (ST) and submitted to the Institut Pateur database.

| **Id** | **Phylogroup_from_ST** | **Country** | ***gapA*** | ***infB*** | ***mdh*** | ***pgi*** | ***phoE*** | ***rpoB*** | ***tonB*** | **ST (MLST)** |
| --- | --- | --- | --- | --- | --- | --- | --- | --- | --- | --- |
| 6358 | Kp1 (7 loci) | Senegal | 2 | 3 | 1 | 1 | 10 | 1 | 19 | 13 |
| 386 | Kp1 (7 loci) | Senegal | 2 | 1 | 1 | 1 | 4 | 4 | 4 | 17 |
| 388 | Kp1 (7 loci) | Senegal | 2 | 1 | 1 | 1 | 4 | 4 | 4 | 17 |
| 6234 | Kp1 (7 loci) | Senegal | 2 | 1 | 1 | 1 | 4 | 4 | 4 | 17 |
| 6246 | Kp1 (7 loci) | Senegal | 2 | 1 | 1 | 1 | 4 | 4 | 4 | 17 |
| 6250 | Kp1 (7 loci) | Senegal | 2 | 1 | 1 | 1 | 4 | 4 | 4 | 17 |
| 28320 |  | Senegal | 2 | 1 | 1 | 1 | 4 | 4 | 4 | 17 |
| 6345 | Kp1 (7 loci) | Senegal | 2 | 3 | 2 | 2 | 6 | 4 | 4 | 29 |
| 6249 | Kp1 (7 loci) | Senegal | 2 | 1 | 2 | 1 | 7 | 1 | 7 | 36 |
| 6289 | Kp1 (7 loci) | Senegal | 2 | 1 | 2 | 1 | 7 | 1 | 7 | 36 |
| 7217 | Kp1 (7 loci) | Senegal | 2 | 1 | 2 | 1 | 7 | 1 | 7 | 36 |
| 28938 |  | Senegal | 2 | 1 | 2 | 1 | 7 | 1 | 7 | 36 |
| 387 | Kp1 (7 loci) | Senegal | 2 | 9 | 2 | 1 | 13 | 1 | 16 | 37 |
| 390 | Kp1 (7 loci) | Senegal | 2 | 9 | 2 | 1 | 13 | 1 | 16 | 37 |
| 6311 | Kp1 (7 loci) | Senegal | 2 | 9 | 2 | 1 | 13 | 1 | 16 | 37 |
| 6362 | Kp1 (7 loci) | Senegal | 2 | 9 | 2 | 1 | 13 | 1 | 16 | 37 |
| 7223 | Kp1 (7 loci) | Senegal | 2 | 1 | 2 | 4 | 9 | 1 | 14 | 39 |
| 6315 | Kp1 (7 loci) | Senegal | 2 | 1 | 1 | 6 | 7 | 1 | 12 | 45 |
| 28423 |  | Senegal | 2 | 1 | 1 | 6 | 7 | 1 | 12 | 45 |
| 6256 | Kp1 (7 loci) | Senegal | 2 | 1 | 5 | 1 | 17 | 4 | 42 | 111 |
| 385 | Kp1 (7 loci) | Senegal | 2 | 20 | 23 | 1 | 1 | 1 | 19 | 129 |
| 389 | Kp1 (7 loci) | Senegal | 2 | 20 | 23 | 1 | 1 | 1 | 19 | 129 |
| 6268 | Kp4 (7 loci) | Senegal | 18 | 22 | 26 | 23 | 31 | 13 | 49 | 138 |
| 6344 | Kp4 (7 loci) | Senegal | 18 | 22 | 26 | 23 | 31 | 13 | 49 | 138 |
| 8275 |  | Senegal | 19 | 23 | 26 | 22 | 32 | 13 | 51 | 141 |
| 8276 |  | Senegal | 19 | 23 | 26 | 22 | 32 | 13 | 51 | 141 |
| 8277 |  | Senegal | 20 | 22 | 27 | 24 | 33 | 20 | 51 | 142 |
| 8278 |  | Senegal | 20 | 22 | 27 | 24 | 33 | 20 | 51 | 142 |
| 8279 |  | Senegal | 20 | 22 | 27 | 24 | 33 | 20 | 51 | 142 |
| 8280 |  | Senegal | 20 | 22 | 27 | 24 | 33 | 20 | 51 | 142 |
| 8281 |  | Senegal | 17 | 19 | 29 | 25 | 35 | 21 | 53 | 143 |
| 8282 |  | Senegal | 17 | 19 | 29 | 25 | 35 | 21 | 53 | 143 |
| 8283 |  | Senegal | 17 | 19 | 28 | 20 | 34 | 18 | 52 | 144 |
| 8284 |  | Senegal | 17 | 19 | 28 | 20 | 34 | 18 | 52 | 144 |
| 6286 | Kp1 (7 loci) | Senegal | 4 | 1 | 32 | 1 | 7 | 4 | 10 | 151 |
| 6266 | Kp1 (7 loci) | Senegal | 2 | 3 | 2 | 1 | 1 | 4 | 56 | 152 |
| 7205 | Kp1 (7 loci) | Senegal | 2 | 3 | 2 | 1 | 1 | 4 | 56 | 152 |
| 6297 | Kp1 (7 loci) | Senegal | 10 | 1 | 1 | 1 | 12 | 1 | 38 | 225 |
| 8298 |  | Senegal | 4 | 1 | 1 | 1 | 7 | 1 | 38 | 242 |
| 6347 | Kp1 (7 loci) | Senegal | 2 | 1 | 44 | 1 | 16 | 4 | 66 | 249 |
| 6282 | Kp1 (7 loci) | Senegal | 3 | 1 | 1 | 1 | 1 | 1 | 43 | 277 |
| 6232 | Kp1 (7 loci) | Senegal | 7 | 1 | 5 | 46 | 1 | 1 | 84 | 281 |
| 6283 | Kp1 (7 loci) | Senegal | 7 | 1 | 5 | 46 | 1 | 1 | 84 | 281 |
| 6313 | Kp1 (7 loci) | Senegal | 2 | 1 | 2 | 1 | 1 | 1 | 4 | 292 |
| 6272 | Kp1 (7 loci) | Senegal | 2 | 1 | 11 | 1 | 1 | 1 | 13 | 337 |
| 6278 | Kp2 (7 loci) | Senegal | 38 | 19 | 53 | 58 | 73 | 21 | 52 | 338 |
| 6273 | Kp1 (7 loci) | Senegal | 2 | 1 | 1 | 1 | 21 | 44 | 9 | 397 |
| 7219 | Kp1 (7 loci) | Senegal | 2 | 1 | 1 | 1 | 21 | 44 | 9 | 397 |
| 7220 | Kp1 (7 loci) | Senegal | 2 | 1 | 1 | 1 | 21 | 44 | 9 | 397 |
| 8328 |  | Senegal | 3 | 3 | 1 | 1 | 1 | 45 | 18 | 407 |
| 6335 | Kp1 (7 loci) | Senegal | 2 | 1 | 11 | 42 | 26 | 4 | 18 | 411 |
| 6287 | Kp1 (7 loci) | Senegal | 2 | 1 | 1 | 4 | 89 | 4 | 4 | 443 |
| 6321 | Kp1 (7 loci) | Senegal | 51 | 1 | 5 | 1 | 9 | 4 | 13 | 491 |
| 6329 | Kp1 (7 loci) | Senegal | 2 | 53 | 3 | 1 | 10 | 4 | 18 | 502 |
| 6314 | Kp1 (7 loci) | Senegal | 2 | 1 | 1 | 1 | 3 | 3 | 18 | 504 |
| 6288 | Kp1 (7 loci) | Senegal | 2 | 1 | 1 | 1 | 8 | 1 | 9 | 514 |
| 6239 | Kp1 (7 loci) | Senegal | 32 | 5 | 1 | 1 | 9 | 4 | 18 | 528 |
| 6275 | Kp1 (7 loci) | Senegal | 2 | 3 | 87 | 1 | 12 | 1 | 26 | 629 |
| 7210 | Kp1 (7 loci) | Senegal | 4 | 3 | 1 | 36 | 9 | 10 | 14 | 661 |
| 6359 | Kp1 (7 loci) | Senegal | 2 | 4 | 2 | 1 | 7 | 1 | 12 | 788 |
| 6341 | Kp4 (7 loci) | Senegal | 18 | 15 | 64 | 59 | 11 | 13 | 51 | 841 |
| 7211 | Kp1 (7 loci) | Senegal | 2 | 1 | 99 | 6 | 1 | 1 | 129 | 867 |
| 6319 | Kp1 (7 loci) | Senegal | 2 | 3 | 2 | 37 | 4 | 1 | 15 | 968 |
| 27725 |  | Senegal | 2 | 3 | 2 | 37 | 4 | 1 | 15 | 968 |
| 1224 | Kp1 (7 loci) | Senegal | 10 | 3 | 2 | 2 | 6 | 4 | 4 | 985 |
| 6355 | Kp1 (7 loci) | Senegal | 10 | 1 | 11 | 1 | 4 | 8 | 43 | 1015 |
| 28375 |  | Senegal | 10 | 1 | 11 | 1 | 4 | 8 | 43 | 1015 |
| 6324 | Kp1 (7 loci) | Senegal | 2 | 3 | 4 | 97 | 12 | 1 | 39 | 1049 |
| 6290 | Kp1 (7 loci) | Senegal | 2 | 1 | 1 | 1 | 3 | 3 | 38 | 1087 |
| 6235 | Kp1 (7 loci) | Senegal | 2 | 79 | 1 | 1 | 7 | 4 | 4 | 1119 |
| 6304 | Kp1 (7 loci) | Senegal | 3 | 5 | 95 | 1 | 16 | 1 | 12 | 1189 |
| 6306 | Kp1 (7 loci) | Senegal | 3 | 5 | 95 | 1 | 16 | 1 | 12 | 1189 |
| 6303 | Kp1 (7 loci) | Senegal | 4 | 1 | 1 | 1 | 12 | 1 | 35 | 1263 |
| 6196 | Kp1 (7 loci) | Senegal | 2 | 5 | 118 | 1 | 179 | 1 | 13 | 1303 |
| 6247 | Mixed: Kp4 (6 loci), Kp3 (1 locus) | Senegal | 48 | 22 | 18 | 59 | 92 | 13 | 51 | 1308 |
| 6322 | Kp1 (7 loci) | Senegal | 2 | 1 | 1 | 1 | 9 | 1 | 4 | 1401 |
| 6327 | Kp1 (7 loci) | Senegal | 2 | 1 | 1 | 1 | 9 | 1 | 4 | 1401 |
| 6332 | Kp1 (7 loci) | Senegal | 2 | 1 | 2 | 1 | 10 | 1 | 112 | 1419 |
| 6350 | Kp1 (7 loci) | Senegal | 2 | 1 | 2 | 1 | 10 | 1 | 112 | 1419 |
| 28239 |  | Senegal | 2 | 1 | 2 | 1 | 10 | 1 | 112 | 1419 |
| 7221 | Kp1 (7 loci) | Senegal | 2 | 1 | 11 | 1 | 9 | 1 | 112 | 1658 |
| 6308 | Kp1 (7 loci) | Senegal | 2 | 1 | 1 | 2 | 89 | 4 | 4 | 1661 |
| 7212 | Kp1 (7 loci) | Senegal | 4 | 1 | 1 | 1 | 3 | 44 | 4 | 1662 |
| 6337 | Mixed: Kp1 (6 loci), Kp2 (1 locus) | Senegal | 2 | 1 | 11 | 20 | 1 | 4 | 13 | 1666 |
| 29003 |  | Senegal | 2 | 1 | 11 | 20 | 1 | 4 | 13 | 1666 |
| 6228 | Kp1 (7 loci) | Senegal | 2 | 5 | 118 | 1 | 10 | 1 | 13 | 1886 |
| 27766 |  | Senegal | 2 | 5 | 118 | 1 | 10 | 1 | 13 | 1886 |
| 6262 | Kp1 (7 loci) | Senegal | 2 | 1 | 65 | 1 | 240 | 4 | 43 | 1962 |
| 7218 | Kp1 (7 loci) | Senegal | 2 | 1 | 65 | 1 | 240 | 4 | 43 | 1962 |
| 6318 | Kp1 (7 loci) | Senegal | 2 | 1 | 1 | 3 | 12 | 1 | 5 | 2108 |
| 6296 | Kp1 (7 loci) | Senegal | 2 | 3 | 6 | 4 | 9 | 1 | 4 | 2141 |
| 28093 |  | Senegal | 2 | 3 | 6 | 4 | 9 | 1 | 4 | 2141 |
| 6293 | Kp4 (7 loci) | Senegal | 18 | 22 | 26 | 64 | 143 | 38 | 49 | 2559 |
| 6328 | Kp1 (7 loci) | Senegal | 2 | 1 | 1 | 2 | 27 | 4 | 4 | 2655 |
| 27510 |  | Senegal | 2 | 1 | 1 | 2 | 27 | 4 | 4 | 2655 |
| 6261 | Kp4 (7 loci) | Senegal | 18 | 22 | 25 | 59 | 291 | 61 | 147 | 2685 |
| 6330 | Kp1 (7 loci) | Senegal | 2 | 3 | 1 | 1 | 7 | 4 | 23 | 2715 |
| 6320 | Kp1 (7 loci) | Senegal | 2 | 1 | 2 | 1 | 9 | 1 | 18 | 2800 |
| 27639 |  | Senegal | 2 | 1 | 2 | 1 | 9 | 1 | 18 | 2800 |
| 6302 | Kp1 (7 loci) | Senegal | 2 | 1 | 2 | 1 | 10 | 1 | 9 | 3030 |
| 6195 | Kp4 (7 loci) | Senegal | 165 | 22 | 228 | 64 | 327 | 92 | 51 | 3107 |
| 6229 | Kp1 (7 loci) | Senegal | 2 | 2 | 1 | 1 | 7 | 1 | 24 | 3108 |
| 6230 | Kp1 (7 loci) | Senegal | 25 | 4 | 1 | 1 | 20 | 1 | 22 | 3109 |
| 6231 | Kp4 (7 loci) | Senegal | 18 | 23 | 85 | 61 | 328 | 105 | 228 | 3110 |
| 6233 | Kp1 (7 loci) | Senegal | 2 | 1 | 2 | 1 | 7 | 4 | 5 | 3111 |
| 28130 |  | Senegal | 2 | 1 | 2 | 1 | 7 | 4 | 5 | 3111 |
| 6237 | Mixed: Kp4 (6 loci), Kp1 (1 locus) | Senegal | 18 | 22 | 55 | 89 | 11 | 41 | 4 | 3112 |
| 6240 | Kp1 (7 loci) | Senegal | 31 | 1 | 1 | 1 | 3 | 1 | 64 | 3113 |
| 6242 | Kp3 (7 loci) | Senegal | 16 | 24 | 36 | 27 | 47 | 22 | 424 | 3114 |
| 6243 | Kp4 (7 loci) | Senegal | 167 | 22 | 18 | 22 | 115 | 37 | 179 | 3115 |
| 6248 | Kp1 (7 loci) | Senegal | 10 | 3 | 1 | 1 | 1 | 10 | 9 | 3116 |
| 6251 | Kp3 (7 loci) | Senegal | 16 | 64 | 229 | 148 | 331 | 22 | 74 | 3117 |
| 6253 | Kp1 (7 loci) | Senegal | 169 | 1 | 2 | 1 | 1 | 7 | 4 | 3118 |
| 6255 | Kp4 (7 loci) | Senegal | 18 | 22 | 123 | 61 | 333 | 166 | 425 | 3120 |
| 6263 | Kp4 (7 loci) | Senegal | 42 | 15 | 26 | 64 | 169 | 20 | 154 | 3121 |
| 6264 | Kp3 (7 loci) | Senegal | 16 | 18 | 21 | 27 | 57 | 93 | 75 | 3122 |
| 6265 | Kp1 (7 loci) | Senegal | 170 | 3 | 1 | 1 | 1 | 10 | 426 | 3123 |
| 6267 | Kp1 (7 loci) | Senegal | 10 | 3 | 20 | 1 | 1 | 10 | 427 | 3124 |
| 6279 | Kp1 (7 loci) | Senegal | 2 | 1 | 1 | 1 | 7 | 1 | 89 | 3157 |
| 6258 | Kp4 (7 loci) | Senegal | 18 | 23 | 56 | 22 | 269 | 13 | 51 | 3257 |
| 6259 | Mixed: Kp1 (6 loci), Kp4 (1 locus) | Senegal | 2 | 3 | 2 | 1 | 145 | 10 | 427 | 3258 |
| 6269 | Mixed: Kp1 (6 loci), Kp4 (1 locus) | Senegal | 10 | 1 | 20 | 1 | 342 | 4 | 264 | 3260 |
| 6270 | Kp1 (7 loci) | Senegal | 2 | 1 | 2 | 1 | 1 | 4 | 441 | 3261 |
| 6271 | Kp1 (7 loci) | Senegal | 26 | 1 | 1 | 1 | 9 | 10 | 23 | 3262 |
| 6274 | Kp4 (7 loci) | Senegal | 42 | 22 | 208 | 61 | 11 | 38 | 169 | 3263 |
| 6276 | Kp4 (7 loci) | Senegal | 42 | 22 | 56 | 141 | 11 | 180 | 237 | 3264 |
| 6277 | Kp4 (7 loci) | Senegal | 18 | 22 | 18 | 22 | 11 | 38 | 192 | 3265 |
| 6280 | Kp4 (7 loci) | Senegal | 42 | 15 | 18 | 22 | 269 | 13 | 49 | 3266 |
| 6284 | Kp1 (7 loci) | Senegal | 2 | 1 | 1 | 1 | 7 | 4 | 442 | 3267 |
| 29025 |  | Senegal | 2 | 1 | 1 | 1 | 7 | 4 | 442 | 3267 |
| 6285 | Kp1 (7 loci) | Senegal | 4 | 3 | 3 | 1 | 20 | 25 | 25 | 3268 |
| 27838 |  | Senegal | 4 | 3 | 3 | 1 | 20 | 25 | 25 | 3268 |
| 6291 | Kp4 (7 loci) | Senegal | 18 | 22 | 26 | 208 | 31 | 13 | 51 | 3269 |
| 6292 | Kp4 (7 loci) | Senegal | 18 | 22 | 233 | 88 | 348 | 13 | 154 | 3270 |
| 6294 | Kp1 (7 loci) | Senegal | 2 | 5 | 1 | 3 | 7 | 1 | 36 | 3271 |
| 6295 | Mixed: Kp2 (6 loci), Outgroup (1 locus) | Senegal | 82 | 19 | 79 | 20 | 349 | 21 | 443 | 3272 |
| 6298 | Kp1 (7 loci) | Senegal | 4 | 1 | 1 | 97 | 27 | 7 | 4 | 3273 |
| 6299 | Kp2 (7 loci) | Senegal | 38 | 19 | 92 | 39 | 272 | 63 | 148 | 3274 |
| 6301 | Kp1 (7 loci) | Senegal | 2 | 1 | 1 | 2 | 10 | 1 | 4 | 3275 |
| 28946 |  | Senegal | 2 | 1 | 1 | 2 | 10 | 1 | 4 | 3275 |
| 6305 | Kp4 (7 loci) | Senegal | 18 | 22 | 55 | 16 | 94 | 13 | 192 | 3276 |
| 6307 | Kp1 (7 loci) | Senegal | 4 | 4 | 1 | 1 | 7 | 4 | 9 | 3277 |
| 28553 |  | Senegal | 4 | 4 | 1 | 1 | 7 | 4 | 9 | 3277 |
| 6312 | Kp1 (7 loci) | Senegal | 4 | 1 | 1 | 1 | 9 | 1 | 218 | 3278 |
| 6325 | Kp1 (7 loci) | Senegal | 2 | 1 | 218 | 1 | 9 | 8 | 12 | 3280 |
| 6326 | Kp1 (7 loci) | Senegal | 4 | 1 | 2 | 2 | 1 | 4 | 5 | 3281 |
| 6333 | Kp1 (7 loci) | Senegal | 2 | 9 | 2 | 1 | 351 | 1 | 16 | 3282 |
| 6334 | Kp1 (7 loci) | Senegal | 2 | 9 | 2 | 1 | 351 | 1 | 16 | 3282 |
| 6338 | Kp1 (7 loci) | Senegal | 14 | 3 | 2 | 1 | 9 | 1 | 14 | 3283 |
| 28322 |  | Senegal | 14 | 3 | 2 | 1 | 9 | 1 | 14 | 3283 |
| 6339 | Kp4 (7 loci) | Senegal | 18 | 22 | 234 | 61 | 11 | 128 | 169 | 3284 |
| 6342 | Kp4 (7 loci) | Senegal | 18 | 22 | 56 | 63 | 94 | 13 | 51 | 3285 |
| 6343 | Kp1 (7 loci) | Senegal | 2 | 3 | 2 | 1 | 12 | 1 | 112 | 3286 |
| 6354 | Kp1 (7 loci) | Senegal | 2 | 3 | 2 | 1 | 12 | 1 | 112 | 3286 |
| 6346 | Kp1 (7 loci) | Senegal | 2 | 3 | 2 | 1 | 7 | 4 | 23 | 3287 |
| 27817 |  | Senegal | 2 | 3 | 2 | 1 | 7 | 4 | 23 | 3287 |
| 6348 | Kp4 (7 loci) | Senegal | 42 | 22 | 26 | 96 | 352 | 13 | 444 | 3288 |
| 6349 | Mixed: Kp4 (6 loci), Kp4/Kp1 (1 locus) | Senegal | 18 | 22 | 55 | 85 | 127 | 13 | 99 | 3289 |
| 6352 | Kp1 (7 loci) | Senegal | 2 | 1 | 2 | 26 | 7 | 4 | 9 | 3290 |
| 27913 |  | Senegal | 2 | 1 | 2 | 26 | 7 | 4 | 9 | 3290 |
| 6353 | Doubtful: Kp6 (3), Kp3 (3), Outgroup (1) | Senegal | 174 | 117 | 214 | 186 | 304 | 155 | 445 | 3291 |
| 6356 | Kp4 (7 loci) | Senegal | 42 | 22 | 191 | 61 | 11 | 105 | 51 | 3292 |
| 6360 | Kp1 (7 loci) | Senegal | 2 | 1 | 5 | 1 | 10 | 4 | 13 | 3293 |
| 27406 |  | Senegal | 2 | 1 | 5 | 1 | 10 | 4 | 13 | 3293 |
| 6361 | Kp1 (7 loci) | Senegal | 7 | 1 | 1 | 1 | 1 | 4 | 12 | 3294 |
| 6363 | Kp1 (7 loci) | Senegal | 4 | 1 | 11 | 1 | 21 | 1 | 446 | 3295 |
| 6364 | Kp3 (7 loci) | Senegal | 16 | 137 | 21 | 154 | 353 | 22 | 74 | 3296 |
| 6365 | Kp1 (7 loci) | Senegal | 4 | 4 | 1 | 1 | 7 | 4 | 46 | 3297 |
| 28769 |  | Senegal | 4 | 4 | 1 | 1 | 7 | 4 | 46 | 3297 |
| 7206 | Kp1 (7 loci) | Senegal | 2 | 1 | 1 | 4 | 89 | 27 | 4 | 3337 |
| 7208 | Kp1 (7 loci) | Senegal | 2 | 1 | 62 | 97 | 10 | 1 | 91 | 3338 |
| 7209 | Kp1 (7 loci) | Senegal | 2 | 3 | 15 | 2 | 18 | 4 | 4 | 3339 |
| 7213 | Kp1 (7 loci) | Senegal | 4 | 3 | 1 | 1 | 9 | 10 | 9 | 3340 |
| 7214 | Kp1 (7 loci) | Senegal | 2 | 1 | 1 | 1 | 12 | 2 | 4 | 3341 |
| 28059 |  | Senegal | 2 | 1 | 1 | 1 | 12 | 2 | 4 | 3341 |
| 7224 | Kp4 (7 loci) | Senegal | 42 | 22 | 26 | 61 | 193 | 38 | 99 | 3342 |
| 7225 | Kp4 (7 loci) | Senegal | 42 | 22 | 26 | 61 | 193 | 38 | 99 | 3342 |
| 15759 | K. quasivariicola | Senegal | 292 | 228 | 417 | 356 | 531 | 181 | 746 | 5460 |
| 7207 |  | Senegal | 18 | 22 | 26 | 63 | 11 | 13 | 593 | 6458 |
